# MSMCE: A Novel Representation Module for Classification of Raw Mass Spectrometry Data

**DOI:** 10.1101/2025.03.05.641695

**Authors:** Fengyi Zhang, Boyong Gao, Yinchu Wang, Lin Guo, Wei Zhang, Xingchuang Xiong

## Abstract

Mass spectrometry (MS) analysis plays a crucial role in the biomedical field; however, the high dimensionality and complexity of MS data pose significant challenges for feature extraction and classification. Deep learning has become a dominant approach in data analysis, and while some deep learning methods have achieved progress in MS classification, their feature representation capabilities remain limited. Most existing methods rely on single-channel representations, which struggle to effectively capture the structural information within MS data. To address these limitations, we propose a Multi-Channel Embedding Representation Module (MSMCE), which focuses on modeling inter-channel dependencies to generate multi-channel representations of raw MS data. Additionally, we introduce a “residual” connection along the channel dimension, significantly enhancing the classification performance of subsequent models. Experimental results on four public datasets demonstrate that the proposed MSMCE module not only achieves substantial improvements in classification performance but also reduces computational resource consumption and enhances training efficiency, highlighting its effectiveness and generalizability in raw MS data classification.

## 1 Introduction

Mass spectrometry (MS) is a highly versatile and powerful analytical tool used for detecting, characterizing, and quantifying various analytes based on their observed mass-to-charge ratio (m/z) [1]. With technological advancements, the rate of MS data generation and its complexity have increased significantly. The large volume and high dimensionality of MS data pose considerable challenges for data analysis, particularly in tasks such as cancer detection [2]. Effectively and accurately interpreting these complex MS datasets has become a major challenge in MS data analysis.

In the field of MS data classification, traditional machine learning algorithms, such as Support Vector Machines (SVM), logistic regression, Random Forest, and XGBoost, have been widely applied and often serve as performance benchmarks [3–8]. However, when employing these conventional algorithms, data preprocessing is an essential step [10], which includes denoising, baseline correction, peak detection, and peak alignment. These operations help correct issues arising during MS signal acquisition, enhance the robustness of subsequent multivariate analysis, and improve data interpretability [9]. Nevertheless, the high dimensionality of MS data combined with limited sample sizes makes it challenging for traditional methods to effectively meet the demands of data analysis [11]. These methods often fail to fully exploit the latent information within the data, thereby limiting their overall performance.

In recent years, deep learning has emerged as a dominant technology in data analysis, achieving breakthrough advancements in model architectures such as Convolutional Neural Networks (CNNs) [12], Recurrent Neural Networks (RNNs) [13], and Transformers [14]. Its rapid development has surpassed the limitations of traditional machine learning in many aspects. Researchers have begun exploring its applications in the field of mass spectrometry, including disease diagnosis [2,11,15,16], peptide sequencing and identification [17–22], mass spectrometry prediction [23], and mass spectrometry clustering [24–26].

Studies have shown that deep learning can directly capture complex patterns and latent features from raw MS data [22,32,39], demonstrating its great potential in MS classification. However, a core challenge in MS data analysis lies in how to quantitatively and effectively represent MS vectors [22]. To reduce computational costs while maintaining high analytical performance, many deep learning methods aim to embed MS vectors into a lower-dimensional space to improve computational efficiency. Spec2Vec [24], inspired by representation learning techniques from natural language processing, learns embedded representations of MS vectors from large-scale MS data, verifying that the relationships between vector fragments can reflect the structural similarity of different compounds. MS2DeepScore [25] employs a Siamese neural network to learn low-dimensional embeddings of MS vectors for predicting the structural similarity between chemical compounds, which is also applicable to MS clustering. GLERMS [26] enhances the low-dimensional representation of MS vectors through contrastive learning, significantly improving compound identification and MS clustering performance.

In the field of image classification, multi-channel images contain complementary information across different channels, enabling a more comprehensive description of the target compared to single-channel representations [27]. For high-dimensional MS data, multi-channel representations offer greater flexibility than single-channel representations by capturing different levels of feature information across multiple channels. Furthermore, by leveraging inter-channel correlations, multi-channel representations can generate more expressive features. This information integration allows the model to learn deep patterns that are difficult to uncover using single-channel representations, thereby enhancing its ability to recognize complex data patterns and improving classification performance.

Against this background, CNNs have emerged as an ideal tool for multi-channel representation learning due to their powerful feature extraction capabilities. CNNs can capture spatial invariance in input data, such as translation and scale invariance, allowing them to extract latent features from MS vectors and encode them into feature maps. Pooling layers further refine structural features [28]. More importantly, CNNs have the ability to automatically learn feature representations from raw MS data without relying on complex feature engineering [32]. By stacking convolutional layers and applying nonlinear transformations, CNNs dynamically generate multi-channel feature representations, effectively capturing different dimensions and hierarchical structures within MS data. These multi-channel features not only enrich feature representations but also enhance the model’s adaptability to high-dimensional data. Due to CNNs’ ability to integrate feature extraction and classification, they maintain high classification accuracy even when handling noisy MS data or signals with shifts. Studies have demonstrated that CNNs have been widely applied to MS classification tasks, including direct classification of raw MS data, consistently exhibiting outstanding performance [10,29–31].

Therefore, this study aims to design a Multi-Channel Embedding Representation Module that constructs inter-channel dependencies to generate more expressive feature representations, thereby improving both the classification accuracy and generalization ability of models for raw MS data. Specifically, the proposed module first applies a fully connected layer to reduce the dimensionality of high-dimensional MS vectors, preserving key features while reducing computational complexity. Then, through a series of stacked convolutional layers, the dimensionally reduced single-channel MS vectors are embedded into multiple complementary channel representations. This multi-channel embedding approach captures hierarchical feature information within raw MS data and establishes deep correlations between channels, ultimately enhancing the classification performance of subsequent models.

This study systematically explores the application of multi-channel embedding representations in raw MS data classification and evaluates the stability and generalization ability of the proposed method across different datasets. The main contributions of this study are as follows:

1. Multi-Channel Embedding Representation for MS Data: By representing MS vectors as multi-channel embedded features, the proposed method enriches the feature expression of raw data, enabling the model to capture deeper associations between different features during the learning process.
2. End-to-End Training Framework: The study integrates the multi-channel embedding module with a deep classification model to construct a deep learning framework tailored for MS classification tasks. This approach eliminates the need for complex handcrafted feature engineering, allowing the model to automatically learn optimal feature representations in an end-to-end manner.
3. Dimensional Adaptation of Multi-Channel Embedding Representations: Compared to the original single-channel MS vectors, the embedded multi-channel vectors exhibit better adaptability within CNN architectures. Additionally, the dimensional setting of the multi-channel vectors aligns more effectively with sequence models such as Transformers, further enhancing compatibility and performance.

## 2 Materials and Methods

Given a batch of MS data *X* ∈ *R^BxD^*, where *B* represents the batch size and *D* denotes the dimensionality of each MS vector, our objective is to transform *X* into a multi-channel embedding representation that captures both global and local features, facilitating subsequent classification tasks. Figure 1 illustrates the structural design of the proposed MSMCE module.

**Figure 1.**
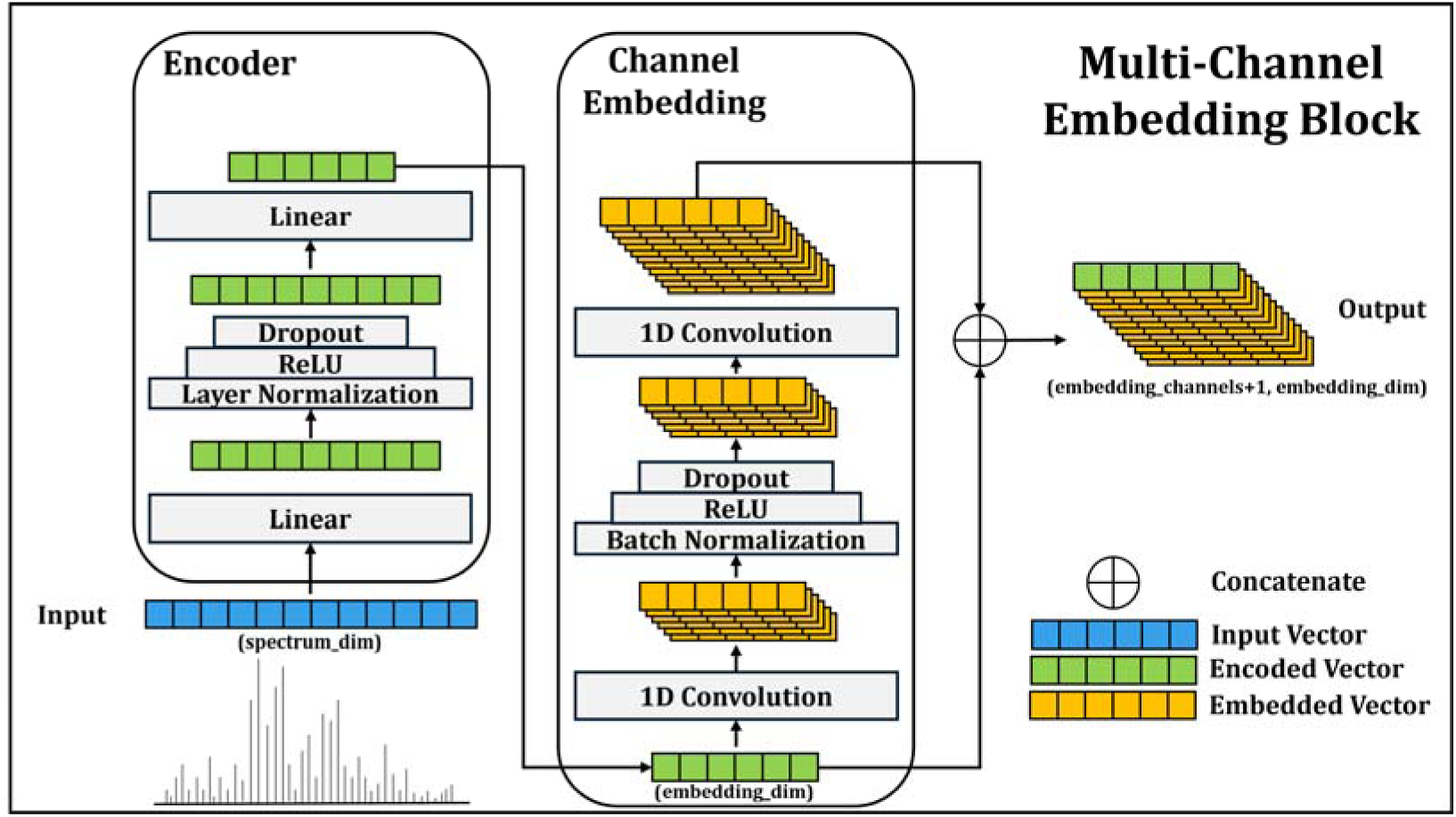
Structural diagram of the Multi-Channel Embedding (MSMCE) module. This module consists of three main components: Encoder, Channel Embedding, and Channel Concat, which are designed to enhance the feature representation capability of mass spectrometry data.

### 2.1 Fully Connected Encoder

The input matrix *X* first passes through a two−layer fully connected network. After the first linear transformation, *X* undergoes layer normalization, ReLU activation, and Dropout, which help prevent overfitting and enhance the model’s generalization ability. This module compresses the dimensionality of the input vectors and extracts global features to generate an initial encoded vector, providing a more refined feature representation for the subsequent channel embedding module. The mathematical formulation is as follows:

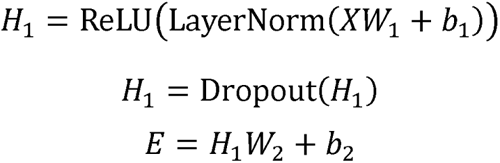

Here, 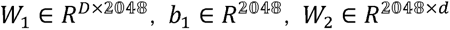, and *b_2_* ∈ *R^d^*. The resulting feature matrix *E* ∈ *R^B×d^* represents the global embedding of the MS data, where *d* is the embedding dimension.

The fully connected encoder aims to reduce the dimensionality of high-dimensional MS vectors through nonlinear transformations, preserving key features while eliminating redundant information. This results in a more compact feature representation, facilitating more effective downstream processing.

### 2.2 Channel Embedding Module

To integrate local features, the encoded feature matrix *E* is reshaped to introduce a channel dimension:

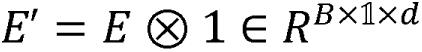

The reshaped tensor is then processed through two consecutive 1D convolutional layers to further extract local features, ultimately producing multi-channel embedded features. After the first convolutional layer, batch normalization, ReLU activation, and a Dropout layer are applied to ensure the model’s generalization capability. The second convolutional layer takes the output of the first layer and extracts deeper local features. Both convolutional layers use the same kernel size to maintain consistency between input and output along the length dimension, ensuring feature scale invariance. The mathematical formulation of the Channel Embedding module is as follows:

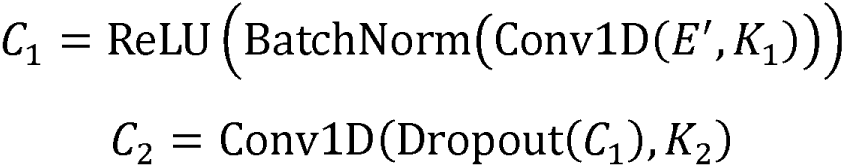

Here, 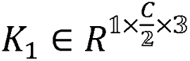 represents the convolution kernel, where *C* is the total number of embedding channels. The output 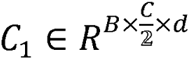 captures local features. The second convolution kernel, 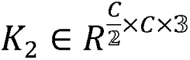, further refines these features, resulting in the final convolutional output 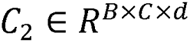, which represents the multi-channel embedded features.

The Channel Embedding module is designed to leverage convolution operations to capture the local structures and spatial relationships within the input data. By transforming feature vectors into multi-channel representations, this module enhances adaptability for downstream classification tasks.

### 2.3 Channel “Residual” Connection

To generate the final output, the original encoded feature *E*’ and the multi−channel embedded feature *C_2_* are concatenated along the channel dimension. This operation is equivalent to applying a residual connection along the channel axis, allowing the embedded features to retain both the global features extracted by the fully connected encoder and the local features captured by the channel embedding module. This fusion enhances the overall feature representation and improves model performance. The channel concatenation can be expressed as:

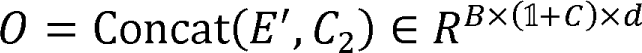

Here, 0 represents the final output tensor of the MSMCE module. The fused feature representation is more diverse and representative, effectively capturing the key feature information within MS data while significantly reducing the dimensionality of the original MS vectors, thereby improving computational efficiency.

The proposed MSMCE module provides an efficient and robust approach for feature extraction from raw MS data. By integrating the global abstraction capability of fully connected layers with the local pattern extraction ability of convolutional layers, this module enables a more comprehensive characterization of the complex features in MS data. In subsequent classification tasks, the MSMCE module demonstrates superior performance, validating its effectiveness and practicality in MS data classification.

**Figure 2.**
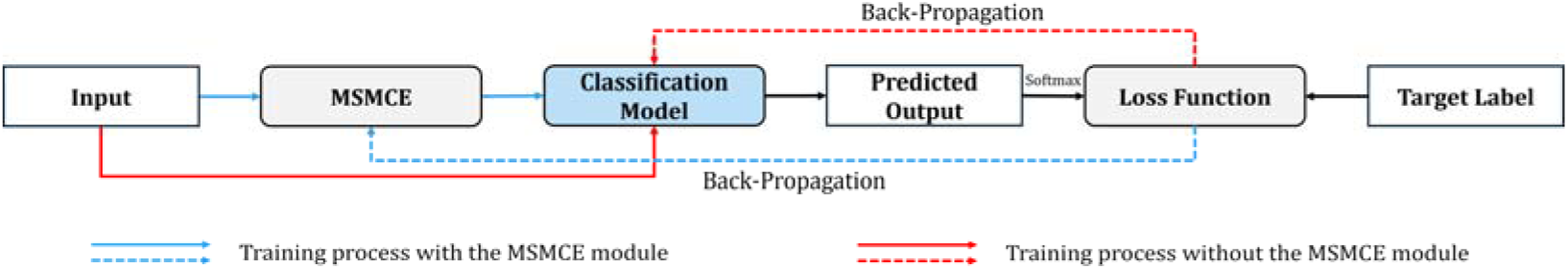
Comparison between the training process with MSMCE and without MSMCE. In the blue path, the input data first passes through the MSMCE module for feature transformation before being fed into the classification model for training. The loss gradient optimizes both the classification model and the MSMCE module. In the red path, the input data is directly fed into the classification model, and the loss gradient is only used to optimize the classification model.

### 2.4 Training Process

To validate the effectiveness of MSMCE in feature representation, we integrate the proposed module with various mainstream deep learning architectures, including CNNs such as ResNet, EfficientNet, and DenseNet, as well as sequence modeling networks like LSTM and Transformer. These models have been widely applied across different tasks, each possessing unique structural characteristics and feature extraction capabilities.

It is important to emphasize that this study does not adopt an ensemble learning strategy. Instead, the multi-channel embedding module serves as a feature representation layer that is directly connected to classification models at the structural level, optimizing feature input for improved classification performance.

As shown in Figure 2, the multi-channel features generated by the MSMCE module are directly fed into the classification model, forming an end-to-end training process. During model training, the multi-channel embedding module and the classification model are jointly optimized, with parameter gradients updated simultaneously under a unified loss function. This approach does not rely on manual feature engineering but instead leverages a deep embedding mechanism to automatically extract and organize feature information from the data, enabling an adaptive representation of the input. The objective is to ensure that different components of the model work collaboratively, maximizing the synergy between the representation module and the classification model, ultimately achieving efficient modeling and classification of raw MS data.

### 2.5 Dataset Description

This study evaluates the proposed method using four publicly available datasets. Table 1 provides detailed information on all datasets, including the mass spectrometry instruments used and the number of classified samples after data processing.

1. Canine Sarcoma Dataset [32]: This dataset contains 1 healthy and 11 sarcoma histology types obtained from 33 annotated ex vivo biopsies. It can be formulated as either a binary classification task or a 12-class classification task [9]. This study analyzes data acquired in positive ion mode.
2. NSCLC Dataset [33]: This dataset includes two major histological subtypes of non-small cell lung cancer (NSCLC)—adenocarcinoma (ADC) and squamous cell carcinoma (SCC)—and can be used as a binary classification task.
3. CRLM Dataset [34]: This dataset consists of colorectal liver metastases (CRLM) tissues and normal liver tissues, serving as a binary classification task.
4. RCC Dataset [35]: This dataset includes samples from renal cell carcinoma (RCC) patients subjects and healthy control subjects, formulated as a binary classification task. This study analyzes data acquired in positive ion mode.

**Table 1.**
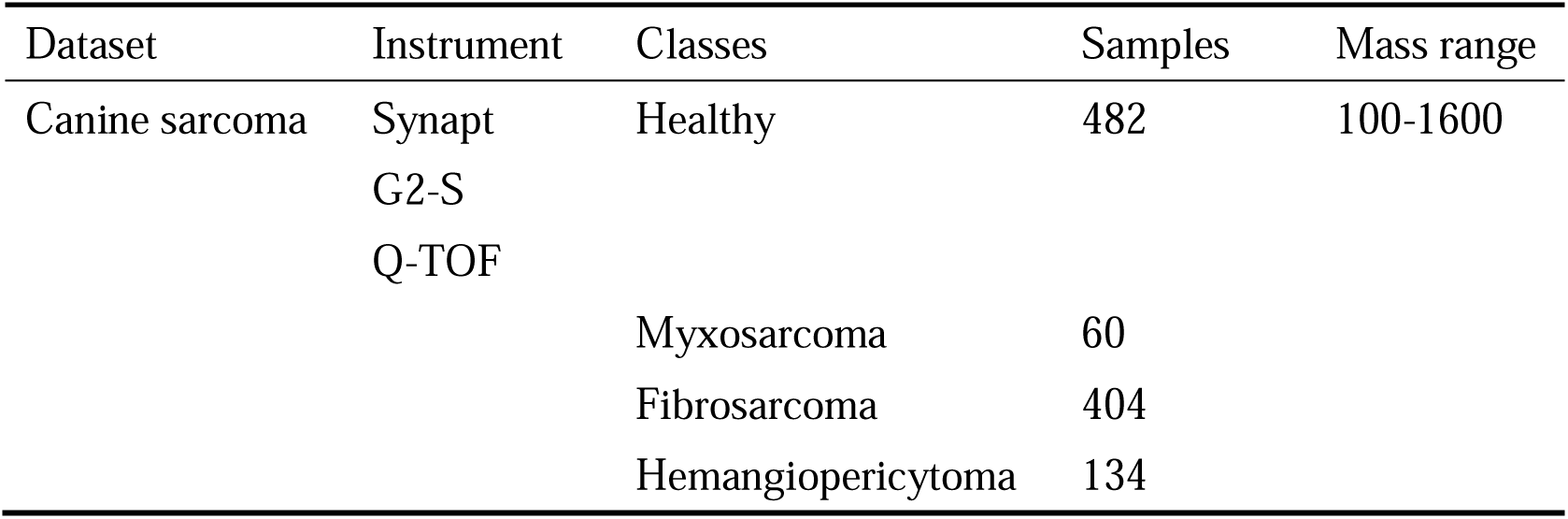
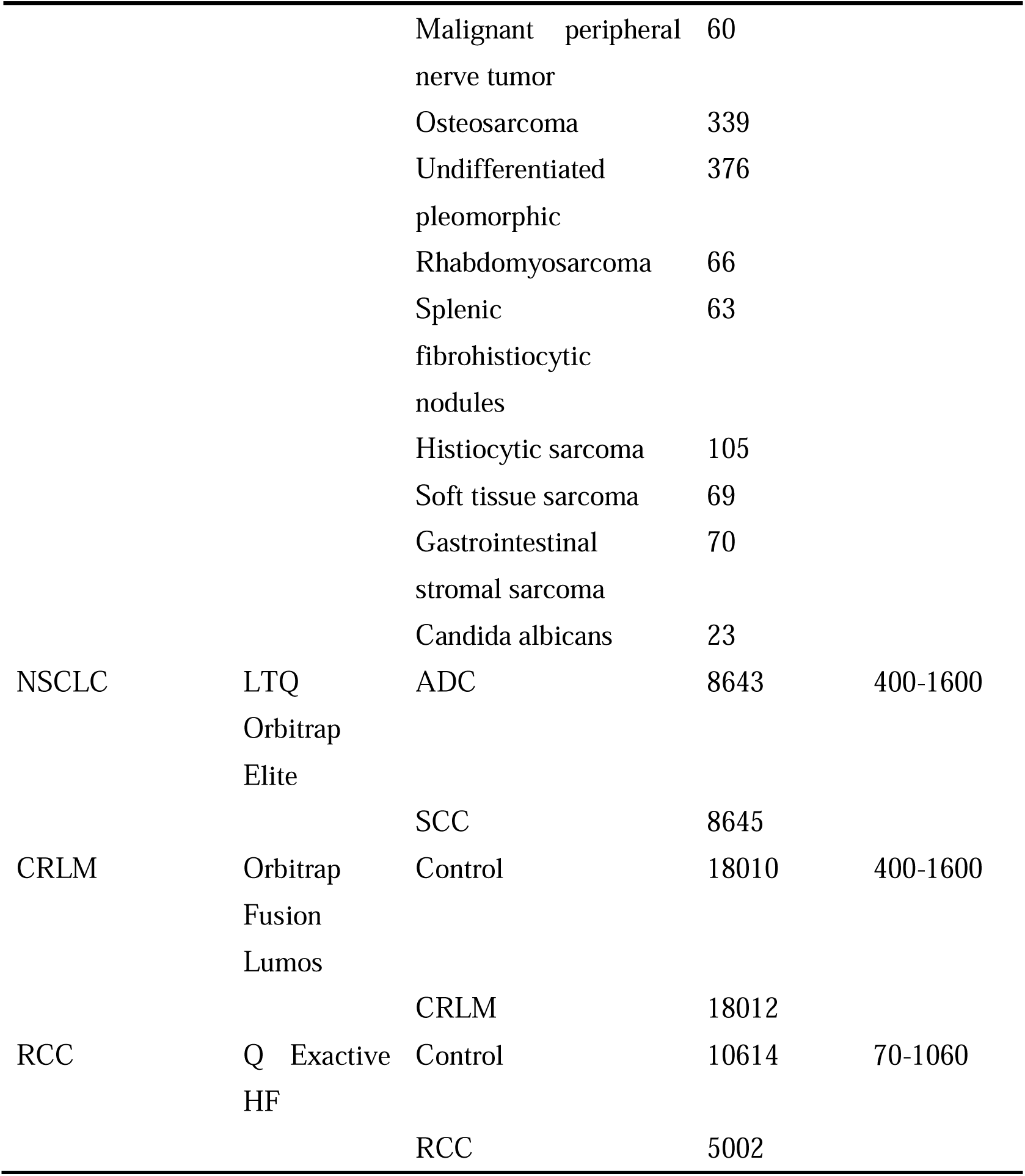
Dataset information.

### 2.6 Data Processing Workflow

In MS data analysis, traditional data processing workflows rely on manual parameter tuning or predefined algorithms for feature engineering, including steps such as peak detection, peak alignment, and feature selection. These processes aim to extract biologically meaningful features while reducing data complexity and noise. However, such methods have inherent limitations: on one hand, preprocessing may lead to the loss of critical information, affecting the accuracy of downstream analyses; on the other hand, despite the development of various data processing workflows, the selection of parameters, quality assessment, and optimal combination of steps lack standardized guidelines. Additionally, traditional feature engineering often requires extensive prior knowledge [38], where handcrafted feature extraction methods must be defined manually. This process is time-consuming, labor-intensive, and difficult to generalize across different tasks.

In recent years, researchers have started exploring strategies for direct classification using raw MS data [2,10,32], reducing the bias introduced by manual feature engineering. These methods typically employ deep learning models that automatically learn key features from raw data. The advantage of this approach is that it preserves more of the original signal information, avoiding the loss of fine-grained features that may occur during traditional preprocessing steps.

This study’s data processing workflow is designed based on recent advancements in the field. Figure 3 illustrates the complete pipeline, from raw mass spectrometry file input to feature matrix generation. For LC-MS datasets, including NSCLC, CRLM, and RCC, we follow the approach described in [2]. The data is first binned with a bin width of 0.1 Da. Next, based on a predefined retention time (RT) binning window—set to 10 seconds—mass spectral vectors within each window are summed and averaged along the feature dimension. For SpiderMass datasets, such as the Canine Sarcoma dataset, we adopt the method outlined in [10]. First, spectra with a total ion count (TIC) below 1 x 10^4^ are filtered out. Then, the spectra are binned with a bin width of 0.1 Da to ensure a consistent data dimension, maintaining comparability across different samples.

**Figure 3.**
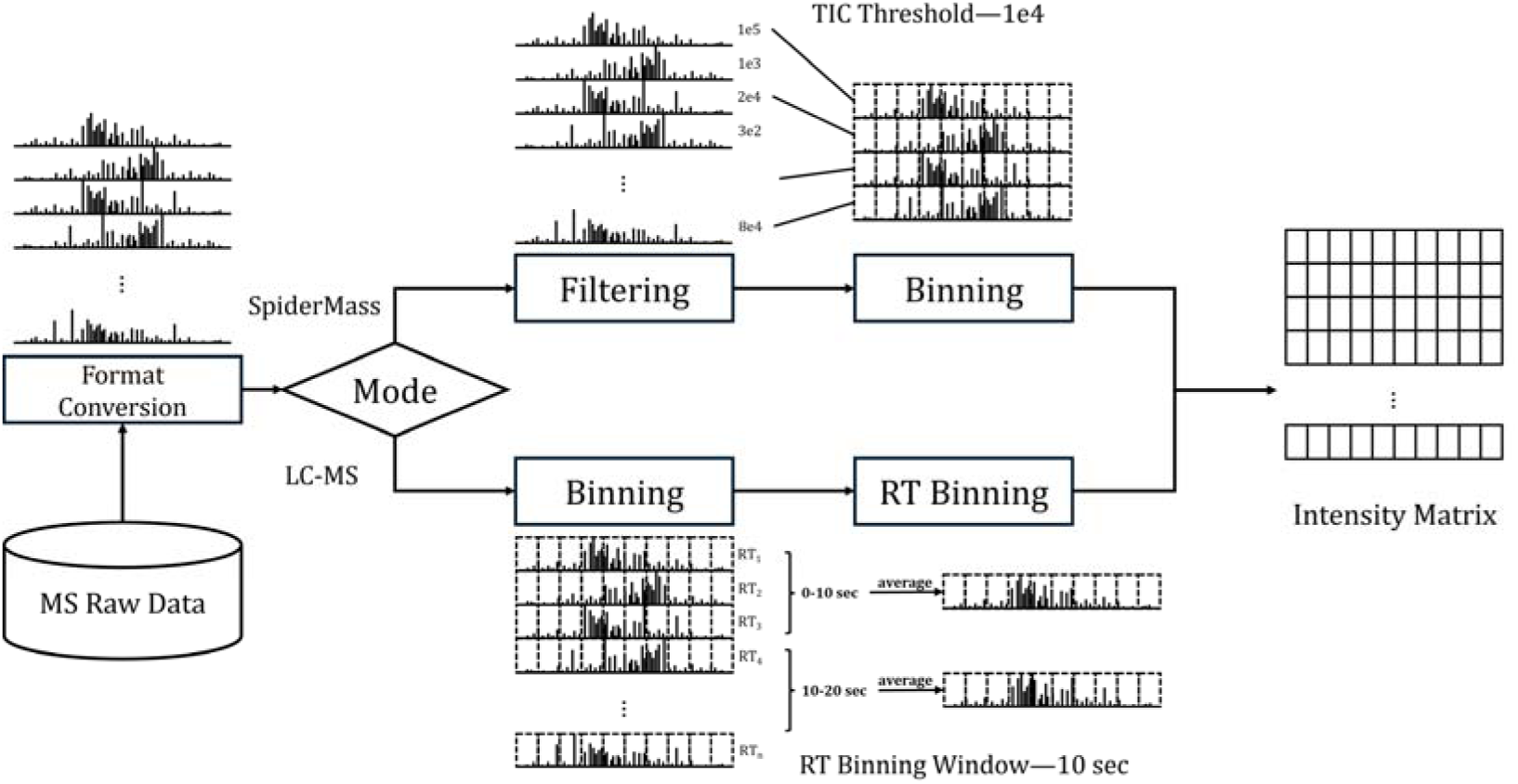
Mass Spectrometry Data Processing Workflow. This figure illustrates the preprocessing pipeline for mass spectrometry data, covering both SpiderMass and LC-MS data processing methods. The final output is an Intensity Matrix, which serves as the input for subsequent analyses.

### 2.7 Experimental Setup and Training Strategy

After the data processing stage, the obtained feature matrix undergoes TIC normalization to reduce the dynamic range of the data and improve feature comparability. TIC normalization adjusts the intensity values of MS data by normalizing the total intensity of each sample to 1.0, thereby eliminating differences in total intensity across samples. This method is widely applied in metabolomics analysis, effectively enhancing sample comparability and improving data consistency.

The normalized data is then split into training, validation, and test sets. To ensure that the class distribution in each subset remains consistent with the original dataset, we employ a stratified sampling strategy, minimizing the impact of class distribution bias on model training and evaluation. To address potential class imbalance in the training data, we compute class weights and apply them to the cross-entropy loss function, enabling class-weighted training to mitigate the influence of imbalance on classification performance. Model parameters are updated using the Adam optimizer, with a learning rate decay scheduler (ReduceLROnPlateau) that automatically adjusts the learning rate when validation performance stagnates, thereby improving model convergence efficiency.

All experiments are trained for 64 epochs. At the end of each epoch, if the validation loss is lower than the historical minimum loss, the minimum loss record is updated, and the current model weights are saved. This ensures that the best model parameters are used for final evaluation on the test set.

## 3 Experiment and Results

To evaluate the effectiveness of the proposed MSMCE module, we conducted a systematic comparison between models embedded with the MSMCE module and their original counterparts without MSMCE across four different datasets. For all experiments, the batch size and mass spectrometry dimensionality were determined based on the characteristics of each dataset. The parameter settings for the MSMCE module remained consistent across all experiments, with the number of channels to embedding set to 256 and the channel dimension set to 1024.

### 3.1 Large-Scale Binary Classification Datasets

The NSCLC, CRLM, and RCC datasets contain a large number of samples, while maintaining simple class structures, as they are all binary classification tasks. Due to their ample sample size and relatively balanced class distribution, these datasets provide an ideal testing environment for evaluating the feature representation capability and classification performance of the proposed models. Conducting experiments on these large-scale datasets aims to assess the adaptability of the MSMCE approach when dealing with high-volume data, particularly in terms of extracting key features, improving classification accuracy, and handling balanced data distributions effectively.

**Table 2.**
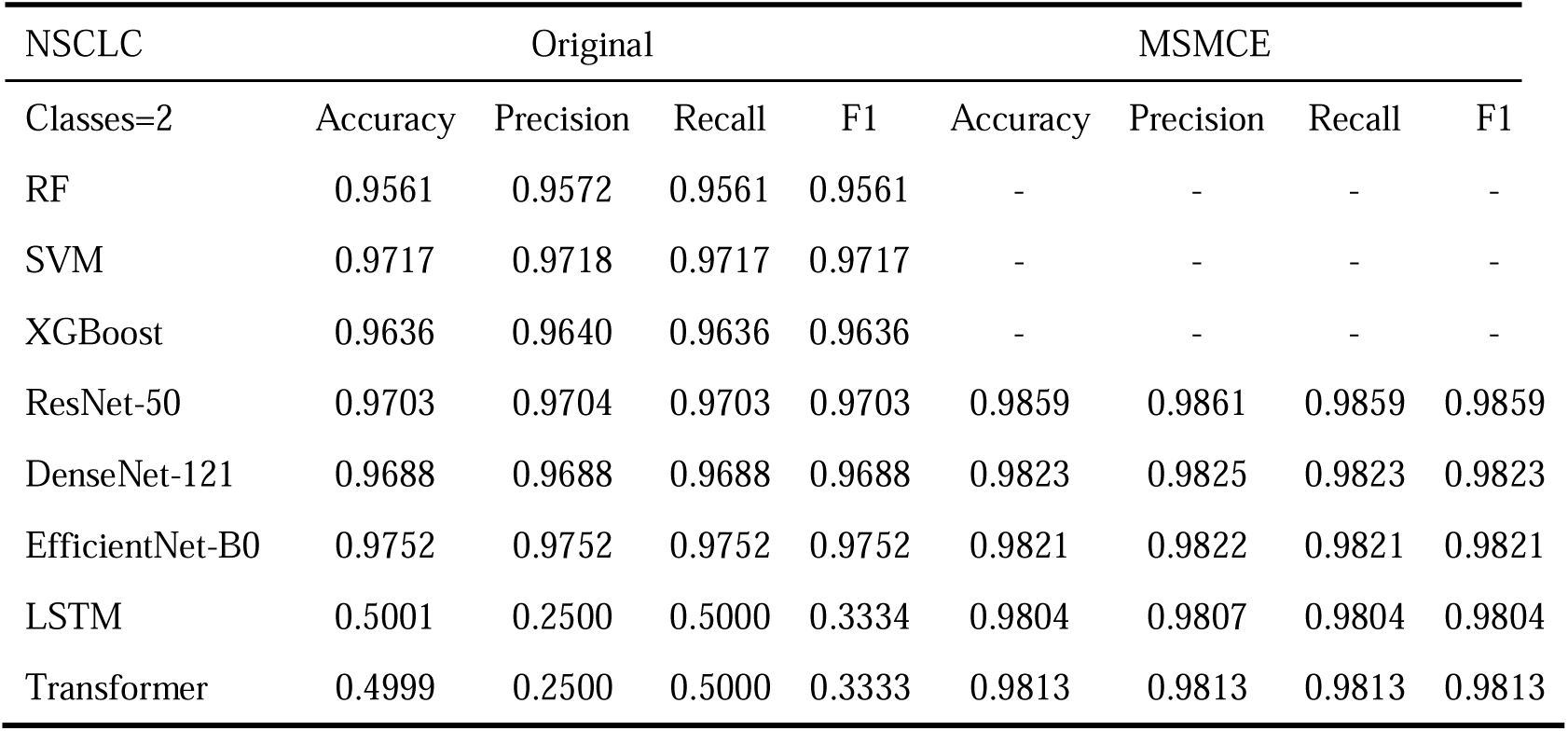
Comparison of Classification Performance of Different Models on the NSCLC Dataset.

**Table 3.** Comparison of Classification Performance of Different Models on the CRLM Dataset.

| CRLM | Original |  |  |  | MSMCE |  |  |  |
| --- | --- | --- | --- | --- | --- | --- | --- | --- |
| Classes=2 | Accuracy | Precision | Recall | F1 | Accuracy | Precision | Recall | F1 |
| RF | 0.8473 | 0.8475 | 0.8473 | 0.8473 | - | - | - | - |
| SVM | 0.8995 | 0.9001 | 0.8995 | 0.8995 | - | - | - | - |
| XGBoost | 0.8812 | 0.8841 | 0.8812 | 0.8810 | - | - | - | - |
| ResNet-50 | 0.8721 | 0.8752 | 0.8720 | 0.8718 | 0.9234 | 0.9254 | 0.9234 | 0.9233 |
| DenseNet-121 | 0.8721 | 0.8737 | 0.8720 | 0.8719 | 0.9287 | 0.9293 | 0.9287 | 0.9286 |
| EfficientNet-B0 | 0.8840 | 0.8870 | 0.8840 | 0.8838 | 0.9203 | 0.9221 | 0.9203 | 0.9203 |
| LSTM | 0.4999 | 0.2499 | 0.5000 | 0.3333 | 0.9126 | 0.9150 | 0.9126 | 0.9124 |
| Transformer | 0.4999 | 0.2499 | 0.5000 | 0.3333 | 0.9215 | 0.9240 | 0.9214 | 0.9213 |

**Table 4.** Comparison of Classification Performance of Different Models on the RCC Dataset.

| RCC | Original |  |  |  | MSMCE |  |  |  |
| --- | --- | --- | --- | --- | --- | --- | --- | --- |
| Classes=2 | Accuracy | Precision | Recall | F1 | Accuracy | Precision | Recall | F1 |
| RF | 0.7741 | 0.8016 | 0.6799 | 0.6950 | - | - | - | - |
| SVM | 0.6746 | 0.6236 | 0.5314 | 0.4851 | - | - | - | - |
| XGBoost | 0.8106 | 0.8337 | 0.7345 | 0.7560 | - | - | - | - |
| ResNet-50 | 0.8033 | 0.7786 | 0.7910 | 0.7837 | 0.8227 | 0.7999 | 0.8046 | 0.8021 |
| DenseNet-121 | 0.7602 | 0.7309 | 0.7363 | 0.7334 | 0.8191 | 0.7957 | 0.8028 | 0.7989 |
| EfficientNet-B0 | 0.8118 | 0.7908 | 0.7782 | 0.7837 | 0.8227 | 0.7997 | 0.8151 | 0.8059 |
| LSTM | 0.6667 | 0.3333 | 0.5000 | 0.4000 | 0.7917 | 0.7674 | 0.7846 | 0.7736 |
| Transformer | 0.6667 | 0.3333 | 0.5000 | 0.4000 | 0.8100 | 0.7868 | 0.8056 | 0.7936 |

In the tables, the Original column represents models without the MSMCE module, while the MSMCE column represents models where MSMCE serves as a feature representation layer. The experimental results demonstrate that deep learning models incorporating MSMCE achieved significant improvements in classification performance across all three datasets, consistently outperforming their original counterparts. As shown in Table 2, on the NSCLC dataset, ResNet-50 with MSMCE achieved an accuracy of 0.9859, representing an increase of 1.608% compared to the original model. As shown in Table 3, on the CRLM dataset, the improvement is even more pronounced. MSMCE-DenseNet-121 achieved an accuracy of 0.9287, a 6.49% improvement over the original DenseNet-121 model. As shown in Table 4, on the RCC dataset, DenseNet-121’s accuracy improved from 0.7602 to 0.8191, an increase of 7.748%, while its F1-score increased from 0.7334 to 0.7989, representing an 8.93% improvement.

The experimental results demonstrate that the MSMCE module effectively leverages the rich sample information in large-scale datasets, capturing complex feature relationships and generating more expressive feature representations. Through the synergistic interaction among multiple channels, this representation enhances the model’s ability to understand the underlying structure and feature patterns in the data, leading to a significant improvement in classification performance.

### 3.2 Small-Scale Multi-Class Classification Dataset

To further validate the applicability of the MSMCE module, experiments were conducted on the Canine Sarcoma dataset. This dataset is characterized by a large number of classes and a limited sample size. It is evaluated for both binary classification and 12-class classification tasks.

**Table 5.**
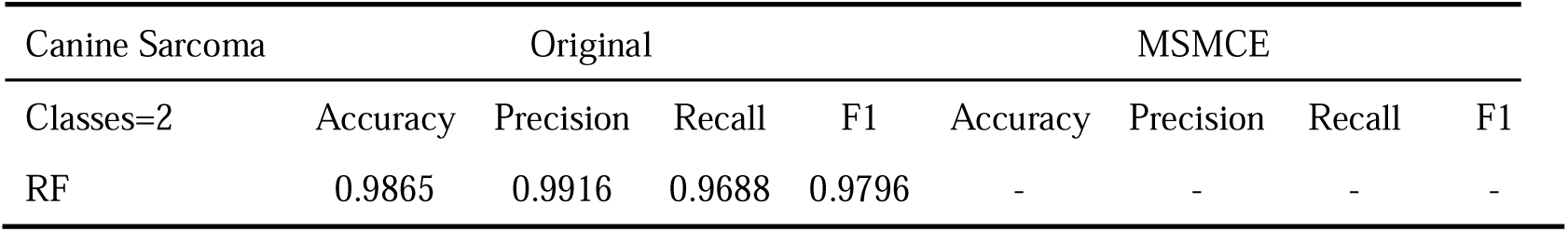
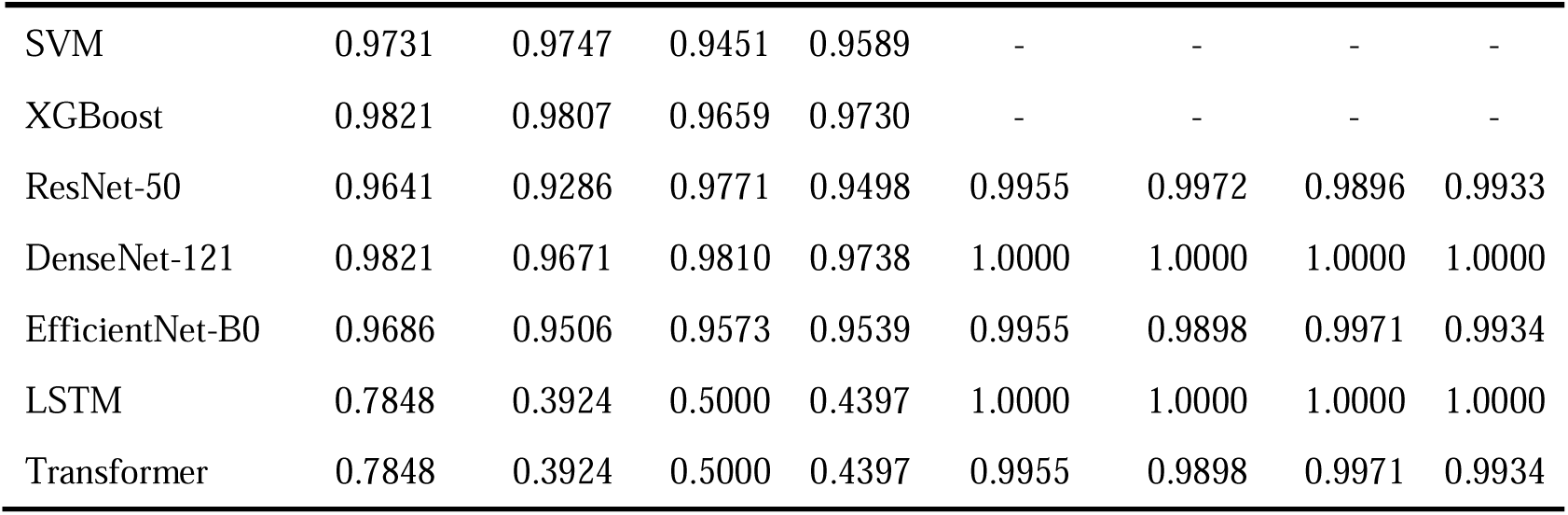
Comparison of Classification Performance of Different Models on the Canine Sarcoma Dataset (Classes = 2).

**Table 6.**
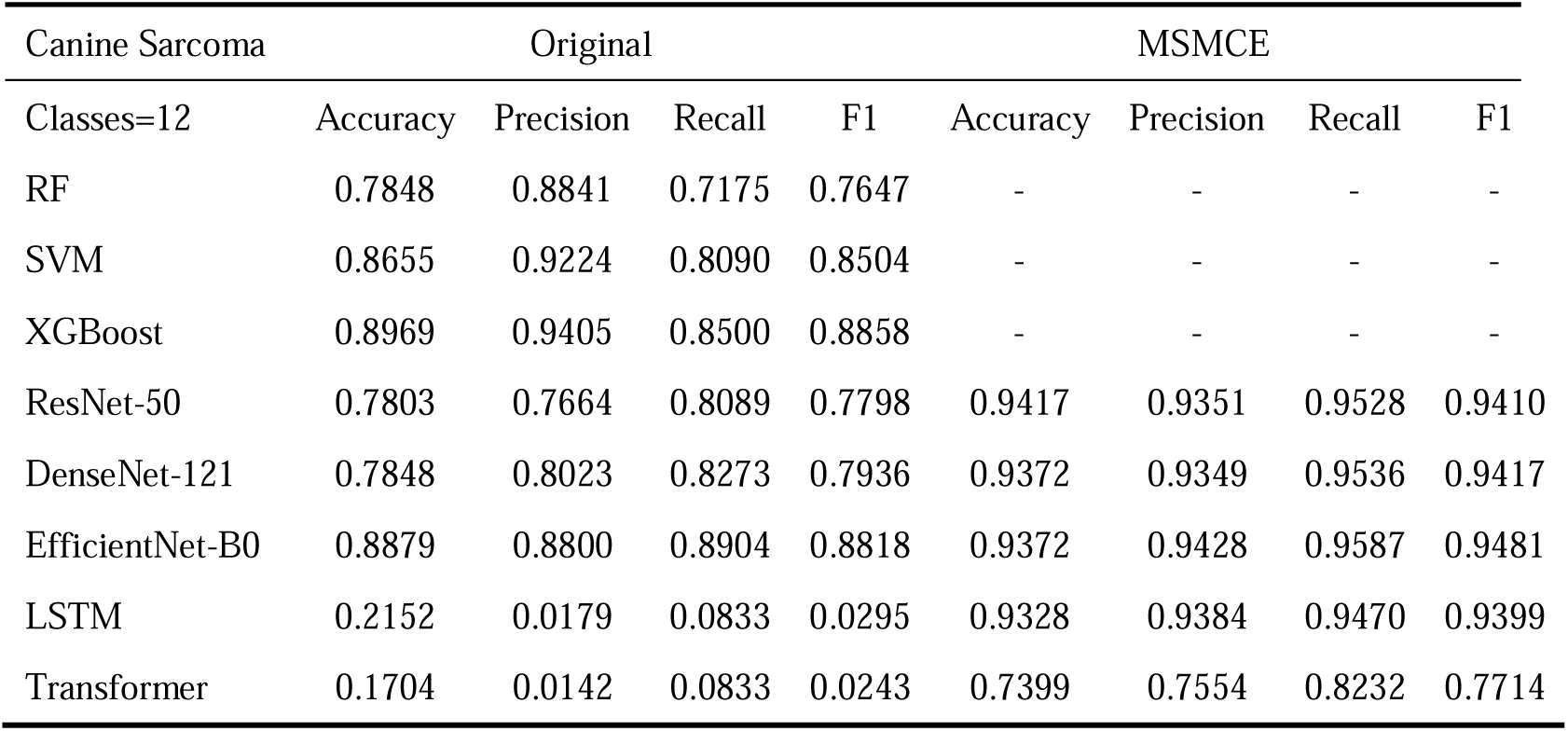
Comparison of Classification Performance of Different Models on the Canine Sarcoma Dataset (Classes = 12).

The experimental results indicate that even on datasets with a limited sample size and a large number of classes, models incorporating the MSMCE module still achieve significant performance improvements. As shown in Table 5, for the binary classification task on the Canine Sarcoma dataset, the accuracy of DenseNet-121 improved from 0.9821 to 1.0000, an increase of 1.79%. For the 12-class classification task, as shown in Table 6, the accuracy of ResNet-50 improved from 0.7803 to 0.9417, representing a 20.68% increase.

These results suggest that the MSMCE module, by constructing multi-channel dependencies, enhances the expressive power of raw mass spectrometry data. Even on datasets with limited samples and multiple classes, models incorporating MSMCE maintain high classification performance. This enhanced feature representation improves the model’s adaptability and robustness when handling highly complex and diverse mass spectrometry data, providing a novel and effective approach for the precise classification of complex diseases.

### 3.3 Ablation Study

To evaluate the actual contribution of the MSMCE module to model performance, we conducted an ablation study on the Canine Sarcoma dataset (Classes = 12). ResNet-50 was selected as the base model, and we progressively introduced the Encoder, Channel Embedding, and Channel Concat to systematically assess their impact on classification performance.

**Table 7.** Ablation Study Results: Impact of Different Module Combinations on ResNet-50 Classification Performance.

| Baseline<br>(ResNet-50) | Encoder | Channel<br>Embedding | Channel<br>Concat | Accuracy | F1 |
| --- | --- | --- | --- | --- | --- |
| ? | - | - | - | 0.7803 | 0.7798 |
| ? | ? | - | - | 0.8072 | 0.7955 |
| ? | ? | ? | - | 0.8834 | 0.8888 |
| ? | ? | ? | ? | 0.9417 | 0.9410 |

**Figure 4.**
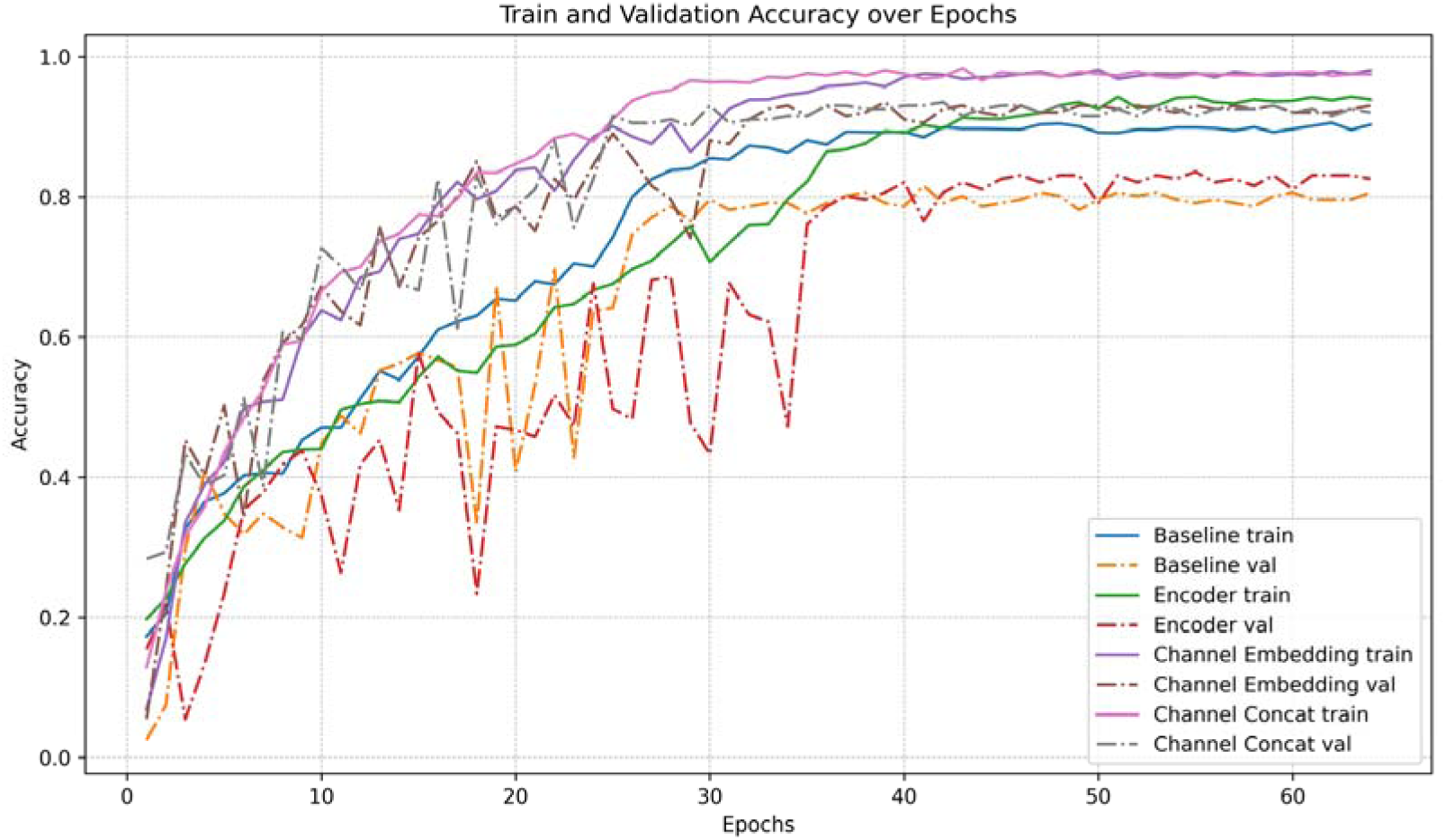
Accuracy Trends During Training and Validation in the Ablation Study. This figure illustrates the accuracy changes on the training and validation sets for ResNet-50 and its variants with progressively introduced MSMCE submodules. The results highlight the impact of different components on model performance. Before introducing the Channel Embedding module, the accuracy curves exhibit significant fluctuations. However, after incorporating Channel Embedding and Channel Concat, the accuracy curves become notably more stable, indicating improved model robustness and convergence.

As shown in Table 7, introducing the Encoder module into the Baseline model leads to a moderate improvement in accuracy and F1-score, indicating that the Encoder plays a role in initial feature extraction. However, the validation accuracy curve for the Encoder exhibits significant fluctuations (see Figure 4), suggesting that the model’s generalization ability is still limited. With the addition of the Channel Embedding module, the model’s performance improves significantly, achieving an accuracy of 0.8834 and an F1-score of 0.8888. This result demonstrates that the Channel Embedding module effectively captures complex relationships between multiple channels, enhancing feature representation. Additionally, the validation performance becomes more stable. Building on this, incorporating the Channel Concat module, which fuses encoded features with channel-embedded features, further optimizes model performance, reaching its best results. This highlights the importance of feature fusion in enhancing classification performance by integrating global and local feature information.

### 3.4 Computational Efficiency Analysis

This study compares the performance of different models in terms of computational complexity, storage requirements, and classification accuracy, while further analyzing the impact of the MSMCE module on computational efficiency. As shown in Figure 5, ResNet-50, DenseNet-121, and EfficientNet-B0 exhibit higher FLOPs (floating point operations) and larger model sizes, indicating greater computational overhead. However, after incorporating the MSMCE module, the FLOPs of all models significantly decrease, and the model sizes are substantially reduced, while classification accuracy improves considerably. These findings suggest that the MSMCE module optimizes computational resource utilization by employing an efficient feature embedding approach, enabling models to achieve superior performance with reduced computational cost.

**Figure 5.**
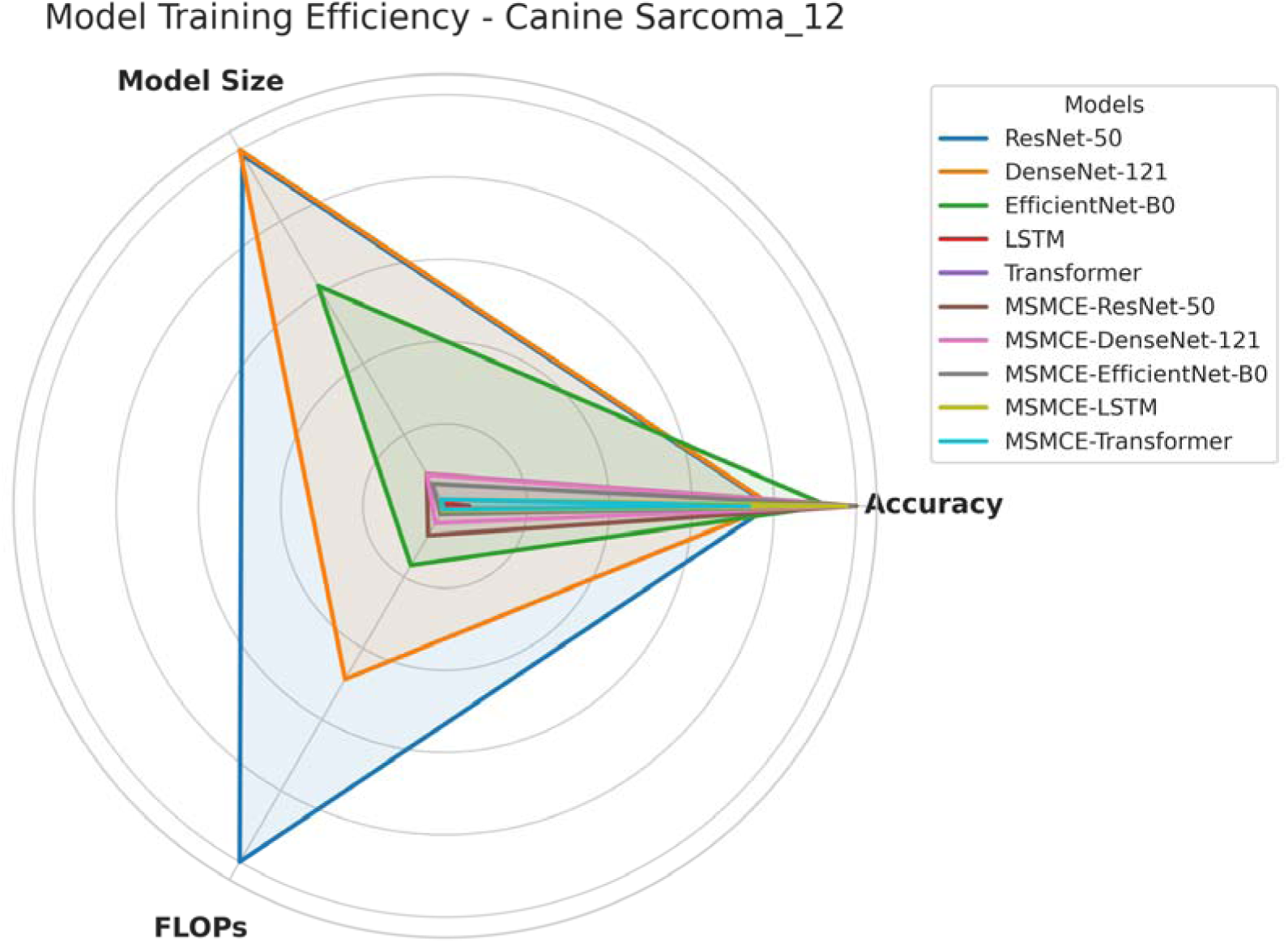
Radar Chart of Model Training Efficiency on the Canine Sarcoma (12-class) Dataset. This figure presents the computational efficiency of ResNet-50, DenseNet-121, EfficientNet-B0, LSTM, and Transformer, along with their corresponding MSMCE-enhanced versions. It can be observed that MSMCE-enhanced models achieve significantly improved classification accuracy while maintaining lower computational costs (FLOPs) and smaller model sizes. Radar charts for training efficiency on other datasets are provided in the supplementary materials.

This performance improvement is attributed to the MSMCE module, which first reduces the dimensionality of high-dimensional data before channel embedding. This significantly decreases the data dimension, ensuring that subsequent convolution operations require less computational effort to capture structural information, thereby reducing unnecessary computational overhead in high-dimensional sparse spaces. Additionally, although LSTM and Transformer models experience an increase in computational demand (higher FLOPs and model size) after incorporating MSMCE, their training stability improves significantly, preventing training crashes.

Notably, while the MSMCE module itself increases the model’s depth, it ultimately enhances computational efficiency. Furthermore, we observe consistent performance improvements across models of different depths, indicating that the benefits of MSMCE can be effectively integrated with various model architectures, making it suitable for both residual and non-residual network structures.

## 4 Discussion

The performance of machine learning methods largely depends on the choice of data representation (or features) used [37], and deep learning is no exception. In MS data analysis, efficiently and accurately representing MS vectors is a core challenge. Representation learning provides an effective solution to this problem, with its fundamental idea being to learn feature representations that enable models to better capture latent information and structures within the data. Traditional MS analysis methods are often limited to single-channel representations, which inherently lack expressive power and struggle to effectively capture the underlying structural information in MS data.

This study proposes a supervised representation learning method based on multi-channel embedding, which converts raw MS vectors into multi-channel embedded representations, significantly enhancing feature expressiveness. Notably, this approach achieves substantial performance improvements even on datasets with numerous classes and limited sample sizes. Compared to traditional feature engineering methods, the multi-channel embedding representation demonstrates greater adaptability, maintaining robust performance when handling new data and tasks. Furthermore, our experimental results confirm that multi-channel embedding not only provides significant advantages in classification accuracy but also plays a crucial role in improving model stability and generalization capability. Additionally, ablation studies validate the contributions of key components, including Encoder, Channel Embedding, and Channel Concat, further demonstrating the critical role of the multi-channel embedding module in enhancing feature representation and classification performance.

The proposed multi-channel embedding representation learning framework provides a novel and efficient deep feature representation method for MS data, effectively addressing the limitations of traditional feature representation approaches while expanding the potential applications of representation learning in MS data analysis.

### 4.1 Limitations and future directions

Although this study has made significant progress, several aspects require further exploration. First, while multi-channel embedding demonstrates strong adaptability on multi-class, small-sample datasets, its performance may still be limited in extreme cases, such as severely imbalanced class distributions. This is because deep learning models rely heavily on data, and when the sample size is too small, the model may fail to effectively learn the feature representations in the embedding space, leading to training failure. Second, the choice of embedding dimensions and parameter optimization still require dataset-specific tuning. Future research could incorporate adaptive feature selection mechanisms to make feature representations more dynamic and better suited for different datasets. Lastly, a major limitation of deep learning models in clinical decision-making is the lack of well-defined interpretability methods [39]. While multi-channel embedding provides rich feature representations, understanding how embedded features across different channels contribute to MS data classification—and uncovering their biological significance—remains an open research question that warrants further investigation.

Future work can focus on improving the interpretability of the MSMCE module, further uncovering the biological significance of embedded features across different channels to enhance the credibility of the model in clinical applications. Additionally, future studies could evaluate the effectiveness of this module in transfer learning scenarios, exploring its transferability across different datasets and tasks to assess its generalizability in mass spectrometry data analysis.

## 5 Conclusion

This study proposes a supervised representation learning method based on multi-channel embedding (MSMCE), which transforms raw MS vectors into multi-channel embedded representations, significantly enhancing feature expressiveness. Experimental results demonstrate that this method not only improves classification accuracy but also enhances model stability and generalization capability. Additionally, ablation studies validate the critical roles of the Encoder, Channel Embedding, and Channel Concat modules, further proving the effectiveness of the MSMCE module in feature learning. Compared to traditional feature engineering methods, the proposed approach exhibits greater adaptability, maintaining robust performance across different datasets and tasks. Overall, this study provides a novel deep feature representation method for MS data analysis, expanding the application potential of representation learning in the field and offering new technical support for MS data analysis.

## Supporting information

Model Size&FLOPs

Training Efficiency

t-SNE Visualization

## Data availability

The Canine Sarcoma, NSCLC, and CRLM datasets are available through P RIDE, with project IDs PXD010990, PXD000438, and PXD008383, respectively.

The RCC dataset can be accessed via Metabolomics, with project ID ST0017 05.

## Code availability

The source code for this study is available on GitHub at https://github.com/WoodFY/MSMCE.

## Acknowledge

This work was supported by the Science & Technology Fundamental Resources Investigation Program (Grant No. 2022FY101200) and received computational support from the National Institute of Metrology, China.

## Author contributions

F.Z. conducted the data processing, experiments, and wrote the main manuscript, prepared the figures 1-5. X.X. and B.G. provided guidance on research direction and overall project supervision. Y.W., L.G., and W.Z. contributed to the collection of the datasets. All authors reviewed the manuscript.

