## Supplementary material for "MSMCE: A Novel Representation Module for Classification of Raw Mass Spectrometry Data": Training Efficiency

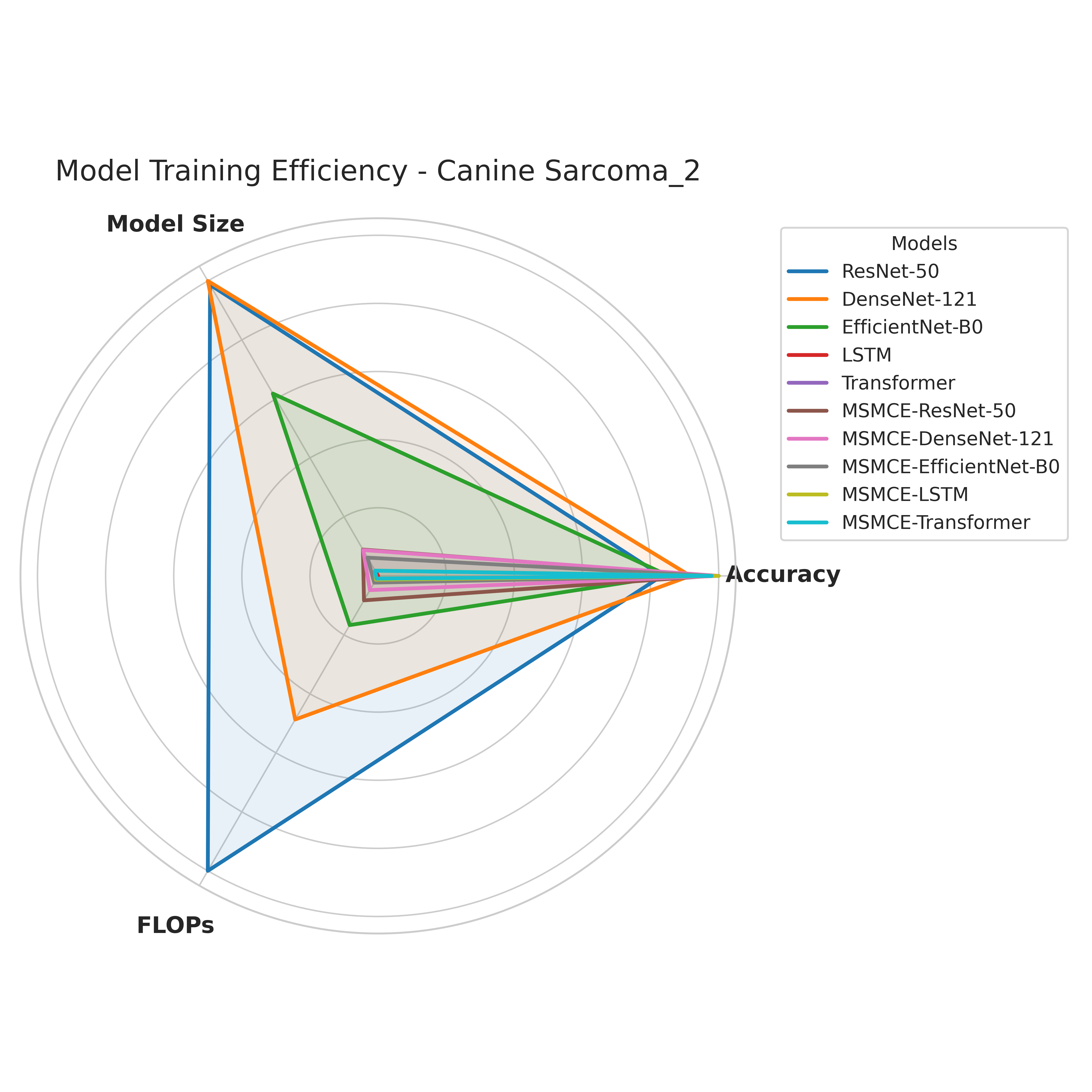

Figure 1 Radar Chart of Model Training Efficiency on the Canine Sarcoma (2-class) Dataset.

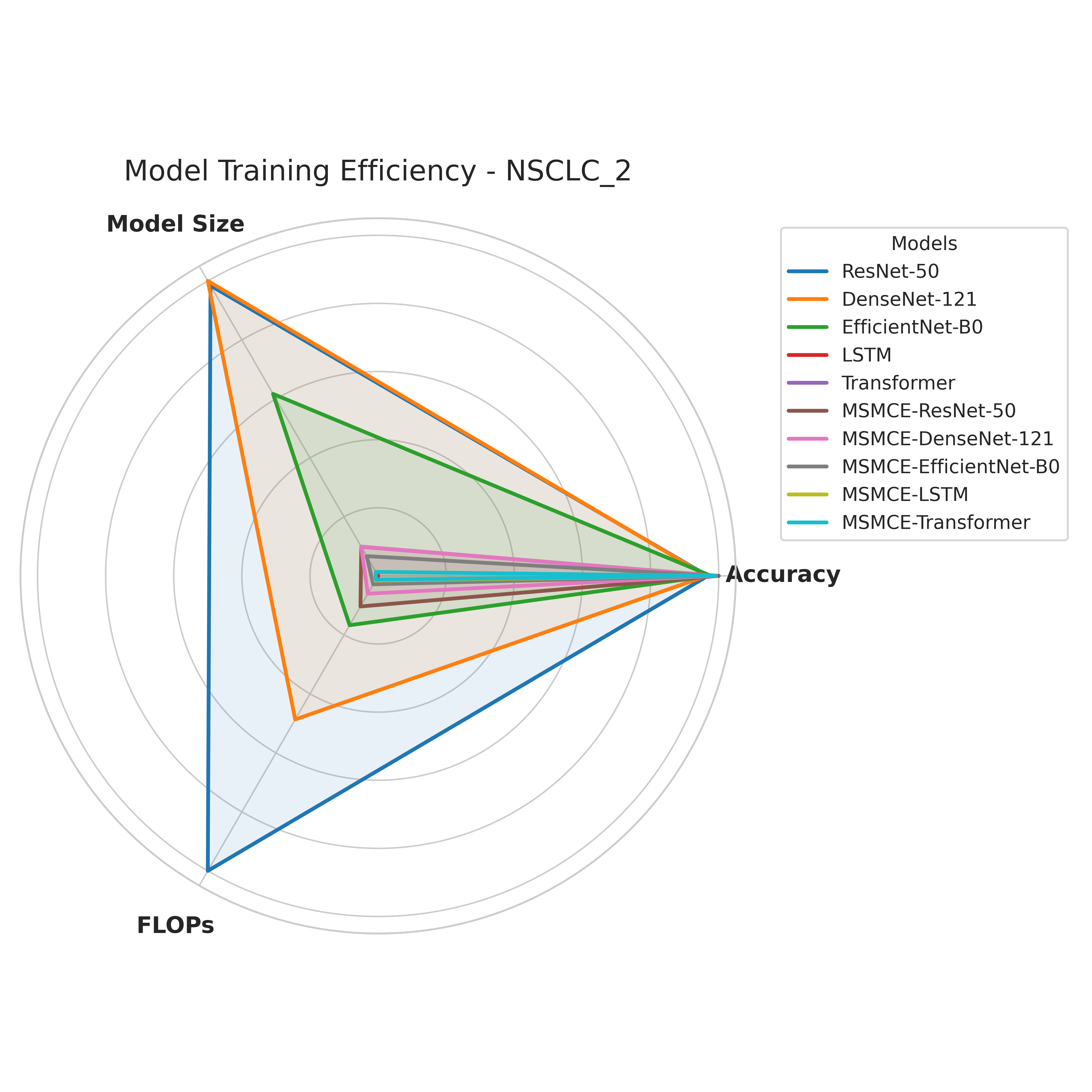

Figure 2 Radar Chart of Model Training Efficiency on the NSCLC (2-class) Dataset.

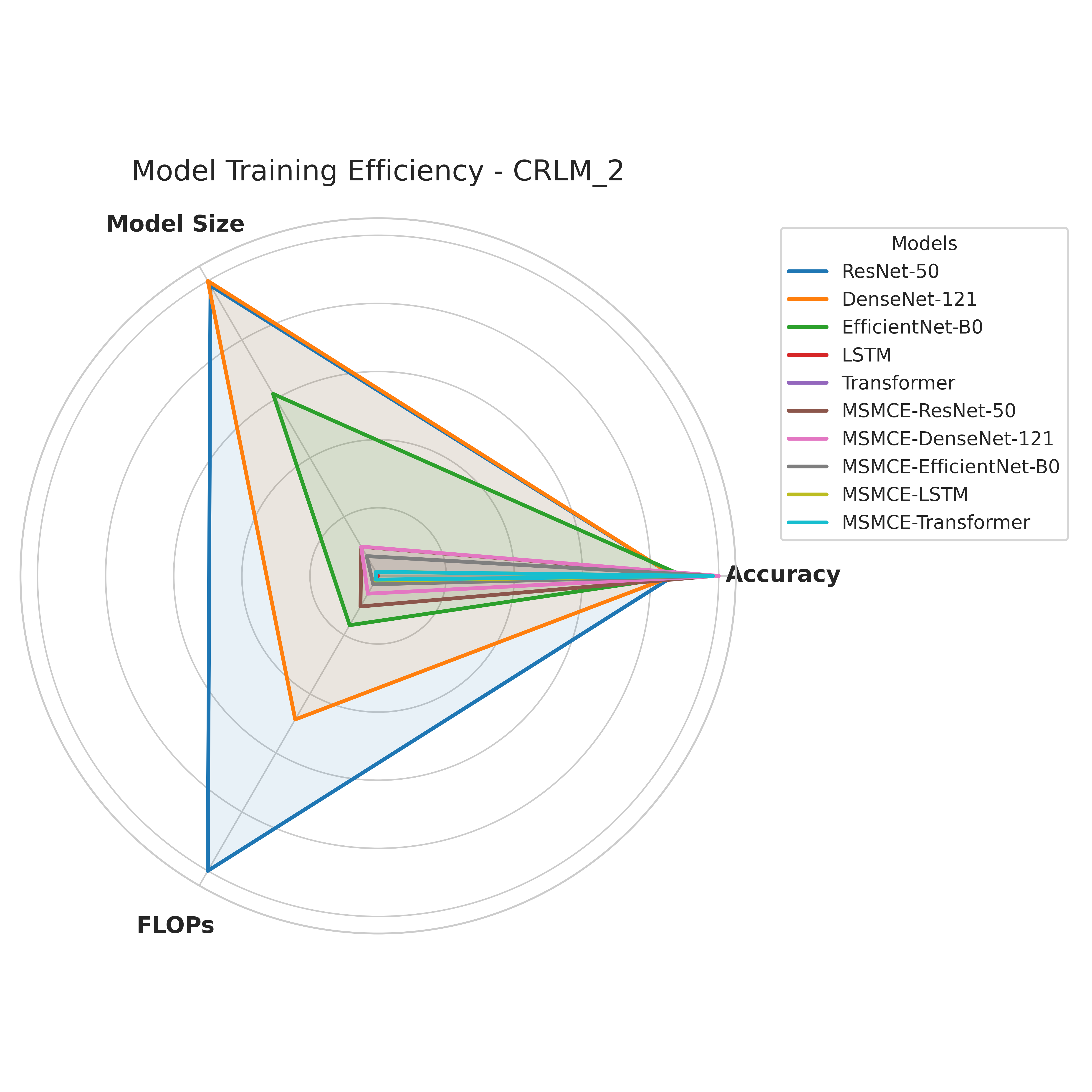

Figure 3 Radar Chart of Model Training Efficiency on the CRLM (2-class) Dataset.

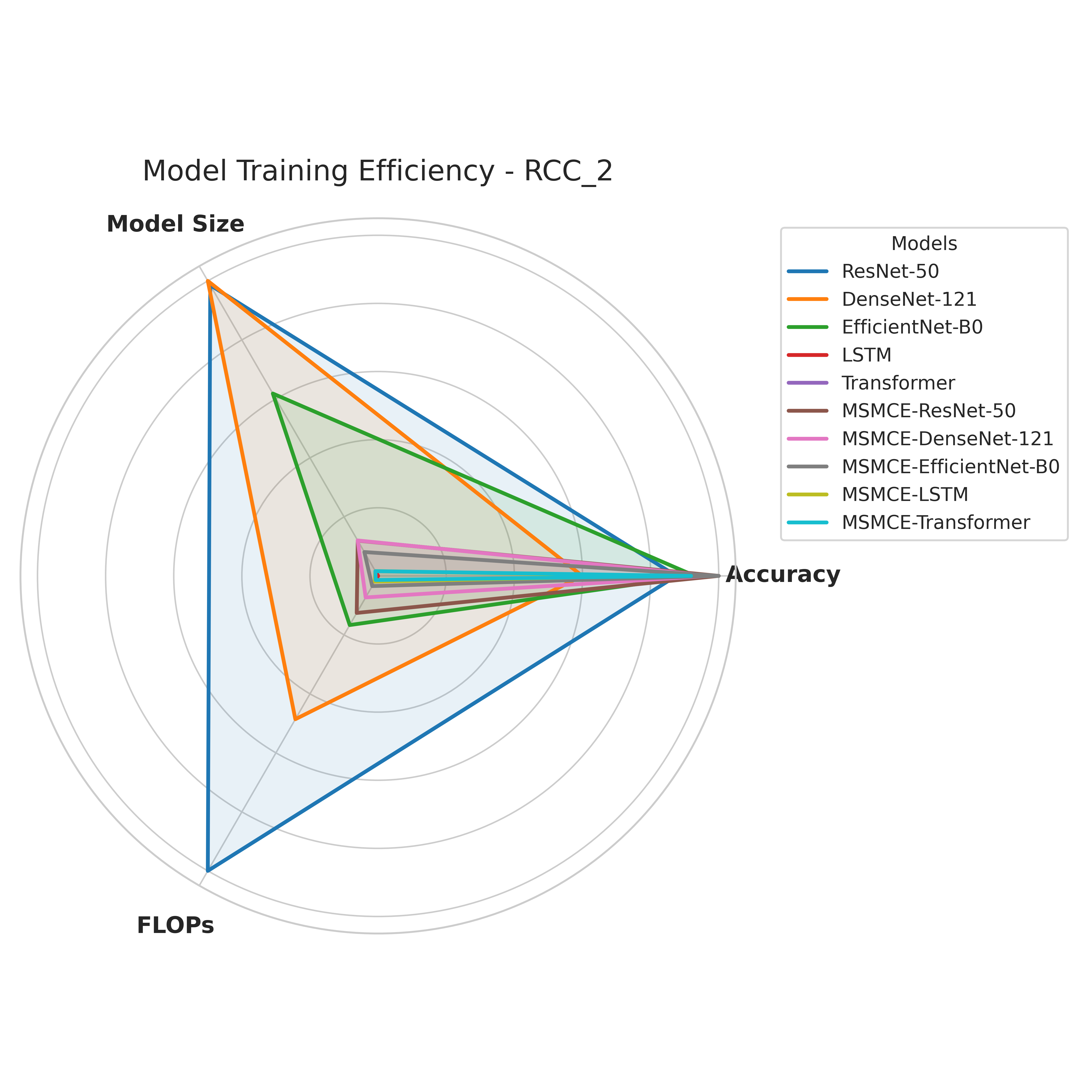

Figure 4 Radar Chart of Model Training Efficiency on the RCC (2-class) Dataset.
