## Supplementary material for "MSMCE: A Novel Representation Module for Classification of Raw Mass Spectrometry Data": t-SNE Visualization

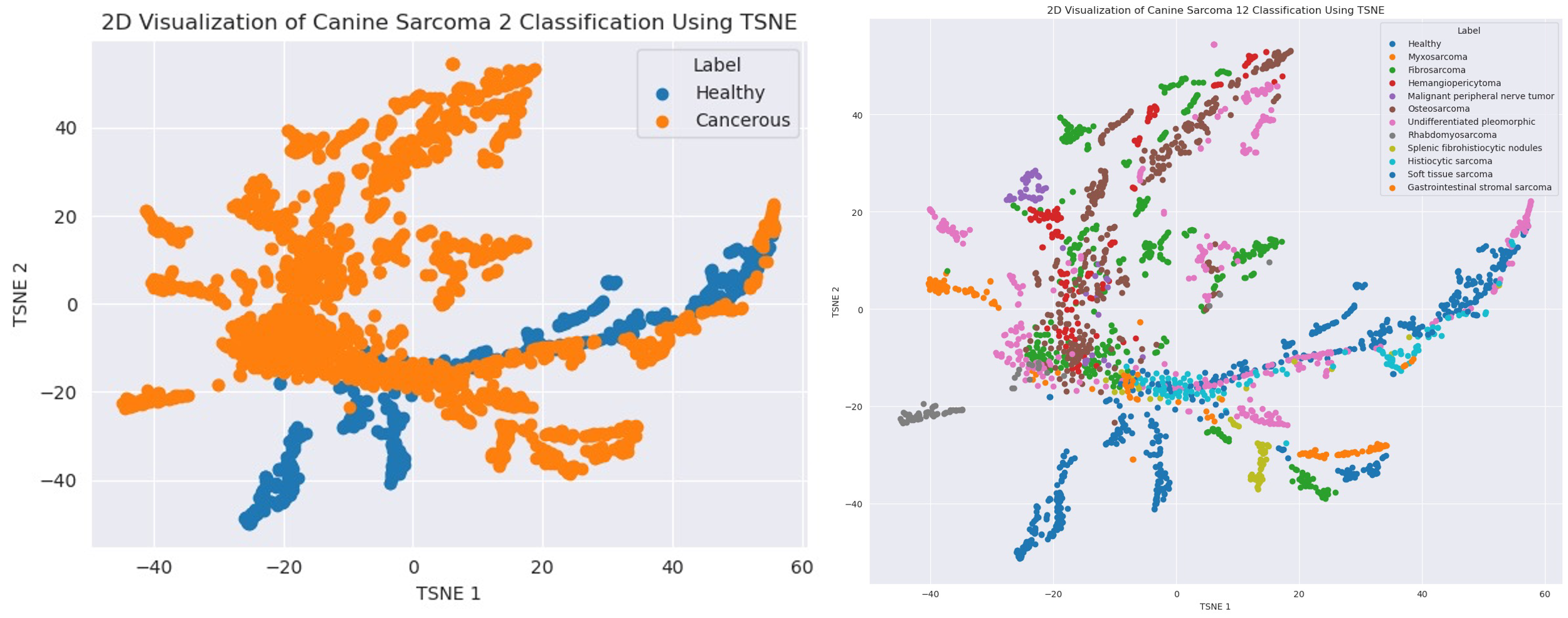

Figure 1 t-SNE Visualizations of Canine Sarcoma Dataset: Binary vs. 12-Class Labels

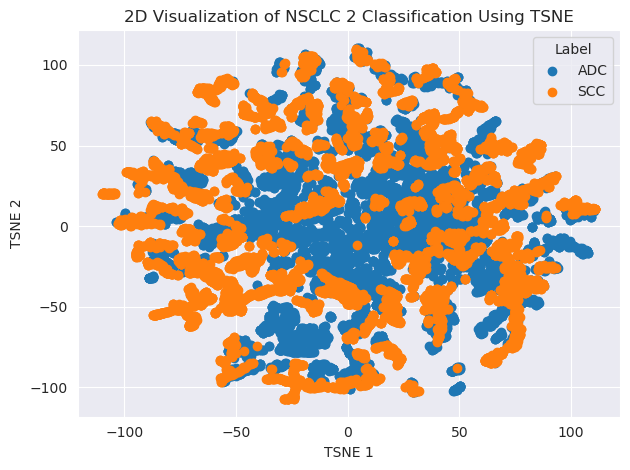

Figure 2 t-SNE Visualizations of NSCLC Dataset: Binary Labels

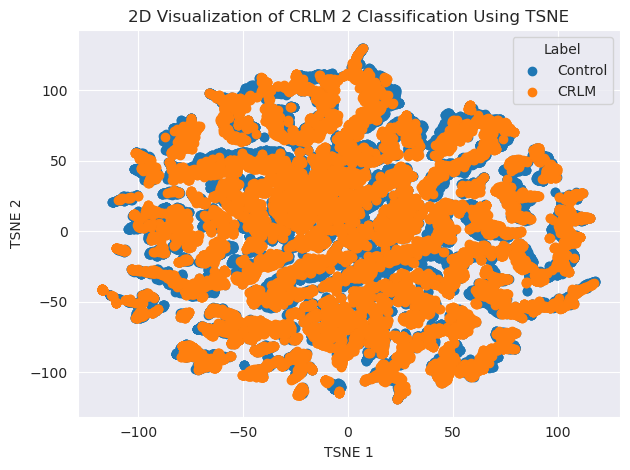

Figure 3 t-SNE Visualizations of CRLM Dataset: Binary Labels

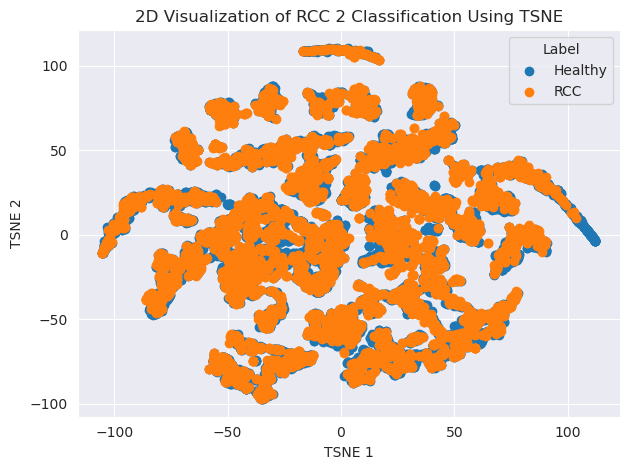

Figure 4 t-SNE Visualizations of RCC Dataset: Binary Labels
